# SpCas9 activity prediction by DeepSpCas9, a deep learning-based model with unparalleled generalization performance

**DOI:** 10.1101/636472

**Authors:** Hui Kwon Kim, Younggwang Kim, Sungtae Lee, Seonwoo Min, Jung Yoon Bae, Jae Woo Choi, Jinman Park, Dongmin Jung, Sungroh Yoon, Hyongbum Henry Kim

## Abstract

We evaluated SpCas9 activities at 12,832 target sequences using a high-throughput approach based on a human cell library containing sgRNA-encoding and target sequence pairs. Deep learning-based training on this large data set of SpCas9-induced indel frequencies led to the development of a SpCas9-activity predicting model named DeepSpCas9. When tested against independently generated data sets (our own and those published by other groups), DeepSpCas9 showed unprecedentedly high generalization performance. DeepSpCas9 is available at http://deepcrispr.info/DeepCas9.

## INTRODUCTION

CRISPR-Cas, a prokaryotic adaptive immune system, has been harnessed for genome editing in various species and cell types, including human cells(1–6). The ability to accurately predict SpCas9 activity is important for applications of genome editing(7–12). So far, several computational models that predict SpCas9 activity have been developed based on data sets of phenotypic changes of gene-edited cells (7,9,11–17) or medium-sized data sets of SpCas9-induced indel frequencies obtained by episomal plasmid-based library-on-library approaches (10,18,19). However, the generalization performances of these models have been limited (20), possibly because the quality and size of the training data sets were not ideal. The majority of models were developed using Cas9 activity data sets generated by phenotypic analysis of gene knockouts (7,9,11–17), which can be biased by the function of the corresponding genes and can include false-negatives in which indels are introduced at the target sequences but do not induce functional knockouts (11); furthermore, for three models, the sizes of the SpCas9-induced indel frequency data sets were just medium-sized (10,18,19).

We recently reported a deep-learning based computational model called DeepCpf1, which predicts AsCpf1 (Cpf1 from *Acidaminococcus sp. BV3L6*) activity with a high generalization performance (21). Our high-throughput evaluation of Cpf1 activity using lentiviral libraries of guide RNA-encoding and target sequence pairs (22) enabled the generation of a large data set of Cpf1-induced indel frequencies, which was used as the training data for DeepCpf1. Although similar paired library-based methods have recently been used to develop computational models that predict the indel sequence patterns generated by Cas9-induced double-strand breaks (23–25), a large data set of Cas9-induced indel frequencies has not been generated, preventing the development of a Cas9-activity predicting computational model with high generalization performance. Here, we developed a high-throughput method for evaluating SpCas9-induced indel frequencies at tens of thousands of target sequences by modifying our previously developed methods for Cpf1 (22), which enabled the development of DeepSpCas9, a deep learning-based model that accurately predicts SpCas9 activities with an unprecedentedly high generalization performance.

## MATERIALS AND METHODS

### Oligonucleotide library design

A pool of 17,840 oligonucleotides was array-synthesized by and purchased from Twist Bioscience (San Francisco, CA). We designed each oligonucleotide to contain a 20-nt guide sequence for the sgRNA, a BsmBI restriction site, a 20-nt barcode sequence (barcode 1), a second BsmBI site, a 15-nt barcode sequence (barcode 2), and the corresponding 30-nt target sequence including a PAM (Supplementary Fig. 1a). Barcode 1 was inserted between the two BsmBI sites to reduce template switching during PCR amplification of the oligonucleotide library (26). Barcode 2, placed upstream of the target sequence, was used to identify each guide RNA and target sequence pair after deep sequencing.

For the target sequences for the oligonucleotide pool, we extracted sequences from the human genome and generated random synthetic sequences without any information about the activity of the corresponding sgRNAs. We first randomly extracted 9,824 target sequences from the GeCKOv1 library (27), excluding those with BsmBI sites in their sequences. From 1,841 target sequences from the coding sequences of three human and six mouse cell surface marker-encoding genes (14) and 2,549 sequences from genes related to resistance against vemurafenib, selumetinib, and 6-thioguanine (7), we obtained 1,804 and 2,484 target sequences, respectively, after excluding sequences containing BsmBI sites. For training the algorithm with guide sequences with extreme GC content, we randomly generated synthetic input sequences containing an NGG PAM with a total length of 30 nt using in-house Python Scripts (Supplementary Code), and subsequently selected 998 input sequences containing guide sequences with extremely low or high GC content (≤ 20% or ≥80%). Additionally, 546 endogenous target sequences from human coding and non-coding genes-of-interest designed for other studies in our lab were included; for this group, five unique barcodes per target sequence were used to yield 5-fold coverage for each target site. Taken together, the set of 17,840 oligonucleotides is composed of 9,824 + (1,804 + 2,484) + 998 + (546 × 5) oligonucleotides.

### Plasmid library preparation

Preparation of the plasmid library containing guide RNA and corresponding target sequence pairs involved a two-step cloning process: Gibson assembly followed by restriction enzyme-induced cutting and ligation (Supplementary Fig. 1). This multistep procedure effectively prevented uncoupling between guide RNA and target sequence pairs during PCR-amplification of the oligonucleotide pool (26). The multistep cloning protocol was adapted and modified from a previously described process (28).

#### Step I: Generation of the initial plasmid library containing guide and target sequence pairs

The Lenti-gRNA-Puro plasmid (Addgene; #84752)(22) was linearized with BsmBI enzyme (NEB, Ipswich, MA) at 55 °C for 6 h. After digestion, the vector was treated with 2 μl of calf intestinal alkaline phosphatase (NEB) at 37 °C for 30 min, then gel-purified using a MEGAquick-spin Total Fragment DNA Purification kit (iNtRON Biotechonology, Seongnam, South Korea).

The oligonucleotide pool was PCR-amplified using Phusion Polymerase (NEB) and primers described in Supplementary Table 4. The amplicons were gel-purified on a 4% agarose gel and assembled with the cut Lenti-gRNA-Puro plasmid described above using an NEBuilder HiFi DNA Assembly kit (NEB). After incubation at 50 °C for 1 h, the assembled products were purified using a MEGAquick-spin Total Fragment DNA Purification kit (iNtRON Biotechnology) and transformed into electrocompetent cells (Lucigen, Middleton, WI) via a MicroPulser (Bio-Rad, Hercules CA). Transformed cells were seeded onto Luria-Bertani (LB) agar plates supplemented with 50 μg/ml carbenicillin and incubated at 37 °C for 16 h. A small fraction (20 μl) of the culture was separately spread to calculate the library coverage; the resulting library coverage ranged from 200× to 220× the initial number of oligonucleotides (i.e., 17,840). Total colonies were harvested and plasmids were extracted using a Plasmid Maxiprep kit (Qiagen, Hilden, Germany).

#### Step II: Insertion of the sgRNA scaffold

The initial plasmid library generated in *Step I* was digested with BsmBI (NEB) for 9 h and treated with 2 μl of calf intestinal alkaline phosphatase (NEB) at 37 °C for 30 min. The digested product was size-selected via 0.8% agarose gel electrophoresis and purified using a MEGAquick-spin Total Fragment DNA Purification kit (iNtRON Biotechnology).

Separately, a synthesized insert fragment containing the sgRNA scaffold was cloned into a TOPO vector (T-blunt vector; Solgent, Daejeon, South Korea). The insert fragment sequence is shown below; the sgRNA scaffold with a poly T sequence is in bold and the BsmBI cut sites are underlined.

<u>CGTCTCT**GTTTT**</u>**AGAGCTAGAAATAGCAAGTTAAAATAAGGCTAGTCCGTTATCAAC TTGAAAAAGTGGCACCGAGTCGGTGCTTT**<u>**TTT**GGGAGACG</u>

Subsequently, the TOPO vector containing the insert fragment was digested with BsmBI (NEB) and the 83-nt insert fragment was gel-purified on a 4% agarose gel. A ligation reaction was performed using 40 ng of this purified insert and 100 ng of the cut initial plasmid library vector described above (Supplementary Fig. 1). Following an overnight incubation at 16 °C, the reaction product was heat inactivated at 65 °C for 10 min and column-purified. The purified product was transformed into electrocompetent cells (Lucigen) via a MicroPulser (Bio-Rad). Transformed cells were seeded onto LB agar plates supplemented with 50 μg/ml carbenicillin and incubated for 16 h at 37 °C. A small fraction of the culture was separately spread on an LB plate with 50 μg/ml carbenicillin to calculate the library coverage. Accordingly, we obtained a final plasmid library coverage of 25~30× the initial number of oligonucleotides (i.e., 17,840). Colonies were harvested and plasmids were extracted using a Plasmid Maxiprep kit (Qiagen).

### Lentivirus production

HEK293T cells (ATCC) were maintained in Dulbecco’s modified Eagle’s Medium (DMEM) supplemented with 10% fetal bovine serum (FBS; Gibco, Waltham MA). For lentivirus production, transfer plasmids containing the gene of interest, psPAX2, and pMD2.G were mixed at a weight ratio of 4:3:1 to yield a total of 20 μg of the plasmid mixture, which was then delivered to 80-90% confluent HEK293T cells using Lipofectamine 2000 (Invitrogen, Carlsbad, CA). At 12 h post-transfection, cells were refreshed with 10 ml of growth medium. The supernatant containing the virus was collected at 36h after the initial transfection, filtered through a Millex-HV 0.45 μm low-protein-binding membrane (Millipore, Darmstadt, Germany), divided into aliquots, and frozen at −80 °C until use. To determine the virus titer, viral aliquots were serially diluted and transduced into HEK293T cells in the presence of 8 μg/ml polybrene. The untransduced cells and serially diluted virus-treated cells were cultured in the presence of 2 μg/ml puromycin or 20 μg/ml blasticidin S (InvivoGen). When almost all of the untransduced cells had died, the number of surviving cells in the virus-treated population was counted to estimate the viral titer as previously described(27).

### Cell library generation

HEK293T cells (9.0 × 10^6^) were seeded into a 150 mm tissue culture dish and incubated overnight. The cells were transduced with the lentiviral library at MOI 0.3 in the presence of 8 μg/ml polybrene and incubated overnight (15 ~ 18 hours). To remove untransduced cells, the cells were cultured in the presence of 2 μg/ml puromycin. To preserve its diversity, the cell library was maintained at a quantity of at least 9.0 × 10^6^ cells throughout the study.

### Cas9 delivery into the cell library

A total of 1.8 × 10^7^ cells (two 150 mm culture dishes with 9.0 × 10^6^ cells per dish) from the cell library were transduced with SpCas9-encoding lentiviral vectors at MOI 5 in DMEM supplemented with 10% FBS and 8 μg/ml polybrene. After overnight incubation (15 ~ 18 hours), the culture medium was replaced with DMEM supplemented with 10% FBS and 20 μg/ml blasticidin S (InvivoGen). The cultures were harvested at 2.9 days after transduction.

### Measurement of indel frequencies at endogenous sites

A total of 124 target sites were selected from the 546 endogenous targets, which are described above in the oligonucleotide library design section, by stratified random sampling (50 targets for DHS regions and 74 targets for non-DHS regions). HEK293T cells were transfected with a mixture of 100 ng of plasmid encoding sgRNA (pRG2; Addgene #104174) and 100 ng of plasmid encoding SpCas9 (pRGEN-Cas9-CMV/T7-Puro-RFP; purchased from Toolgen, Seoul, Korea) at a density of 1.0 × 10^5^ cells/well in a 96-well plate using TransIT-X2 (Mirus Bio, Madison, WI) according to the manufacturer’s instructions. Following an overnight incubation, the culture medium was replaced with DMEM containing 2 μg/ml of puromycin. Cells were harvested and subjected to deep sequencing 3.7 days after the transfection. The average value of indel frequencies from the triplicate studies was used as the representative indel frequency for each target site.

### Deep Sequencing

Genomic DNA was extracted from cell pellets using a Wizard Genomic DNA purification kit (Promega, Fitchburg, WI). For the high-throughput experiment, integrated target sequences were PCR-amplified using 2X Taq PCR Smart mix (Solgent). A total of 576 μg of genomic DNA was used for the first PCR to achieve over 3,000× coverage over the library (assuming 10 μg of genomic DNA for 10^6^ cells)(22). We performed 288 independent 50 μl PCR reactions with an initial genomic DNA concentration of 2 μg per reaction. The PCR products were then combined into a single pool and were purified with a MEGAquick-spin Total Fragment DNA Purification kit (iNtRON Biotechnology) and 20 ng of purified product was subsequently PCR-amplified using primers containing both Illumina adaptor and barcode sequences (Supplementary Table 4). For the cells transfected with SpCas9- and sgRNA-encoding plasmids, we carried out the independent first PCR reactions in a 20 μl reaction volume containing 40 ng of initial genomic DNA template per sample. Then, a second PCR to attach the Illumina adaptor and barcode sequences was conducted in a 20 μl reaction volume using 0.2 μl of the unpurified product from the first PCR. The resulting amplicons were gel-purified and analyzed using HiSeq or MiniSeq (Illumina, San Diego, CA). The primers used for PCRs are shown in Supplementary Table 4.

### Analysis of indel frequencies

Deep sequencing data were analyzed using in-house Python scripts (Supplementary Code), which were modified from previously used code (22). Each guide RNA and target sequence pair was identified using the unique 15-nt barcode sequence located upstream of the target sequence. Insertions or deletions located around the expected cleavage site (i.e., the 8-nt region centered on the middle of the cleavage site) were considered to be Cas9-induced mutations. To exclude the background indel frequencies originating from array-synthesis and PCR amplification procedures, we normalized the observed indel frequency by subtracting the background indel frequency determined in the absence of Cas9 delivery according to the function:

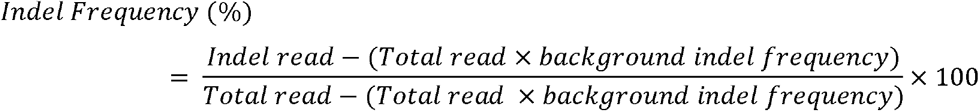

To increase the accuracy of the analysis, deep sequencing data were filtered; target sequences with deep sequencing read counts below 200 and background indel frequencies above 8% were excluded as similarly performed previously(21).

### Calculation of chromatin accessibility

DNase-seq narrow peak data from ENCODE (29) were used to calculate chromatin accessibility as previously described(21). For each target site, 23 bases of the PAM plus protospacer sequence were aligned to the hg19 human reference genome using bowtie(30). Only the target sites that overlapped with DNase-seq narrow peaks were considered as DNase I hypersensitive target sites.

### Convolutional neural network

Convolutional neural networks (CNNs)(31) are one of the most robust deep learning architectures applicable to locally correlated data and have been successfully implemented in DNA sequence-related research such as the prediction of CRISPR-Cpf1 guide RNA activity(21), transcription factor binding affinity(32), and DNA sequence accessibility(33). CNNs consist of three types of layers: a convolutional layer, a pooling layer, and a fully connected layer. In the convolutional layer, various filters are applied to the data, which allows the network to obtain features from local regions rather than the whole. In the pooling layer, several operations (e.g., max, average, etc.) are used to effectively decrease the dimensions and “pool out” useful features from local features extracted from the previous convolutional layer. Convolutional layers and pooling layers are usually interchanged at the initial steps of the CNN, and fully connected layers are constructed in the latter phases. The fully connected layer combines the pooled features by weighted sum and a nonlinear function to get the final function as the solution. Compared to simple multilayer perceptron (MLP) models, CNNs exploit strong heuristics for locally related data. This characteristic has resulted in CNN-based models outperforming the majority of the previously used models in various fields of research.

### Inception module

In CNNs, for each layer, the filter size should be experimentally determined during the model selection phase, as the optimal filter size for the best performance is unknown. In GoogLeNet(34), an inception module was used to overcome this manual process. In an inception module, various sizes of filters are used in one layer as shown in Supplementary Fig. 2. Along with several other techniques, GoogLeNet demonstrated a significant gain in performance compared to the original CNN. Accordingly, we adopted the inception module as our basic module for DeepSpCas9.

### Development of DeepSpCas9

DeepSpCas9 is a deep learning-based computational model for SpCas9 activity prediction. The training data consisted of the HT data set (SpCas9-induced indel frequencies at 12,832 target sequences; Supplementary Table 1) and was used for 10-fold cross validation during the model selection phase. 30 nt-long input sequences, which include 4-bp left neighbor, 20-bp protospacer, 3-bp protospacer adjacent motif, and 3-bp right neighbor sequences, were converted into a four-dimensional binary matrix by one-hot encoding (Supplementary Fig. 2). DeepSpCas9 has one convolutional layer and one pooling layer at the front, and three fully connected layers with a dropout rate of 0.3 in each layer. The adopted convolutional layer includes an inception module with a total of 210 filters (100, 70, and 40 filters at 3, 5, and 7 nts in length, respectively). The pooling layer and three fully connected layers use ReLU activation functions. We tested a total of 324 different models (details in Supplementary Table 5) and selected the model and training epoch that produced the highest validation score calculated using Spearman correlation coefficients between the experimentally measured and predicted activity levels. After selecting the optimal parameters, we used the full training data set with selected parameters to train the final model.

For the development of DeepSpCas9-CA, we fine-tuned DeepSpCas9 using a data subset generated by stratified random sampling of the Endo data set (e.g., Endo-1A) and binary chromatin accessibility information. We added a fully connected layer with 60 units that transformed the binary chromatin accessibility information into a 60-dimensional vector, which enabled the integration of the sequence feature vector and chromatin accessibility information through element-wise multiplication. The regression output layer performs a linear transformation of the outputs and calculates the prediction scores for SpCas9 activity. We applied a dropout rate of 0.3, a mean-squared error, as the objective function, and an Adam optimizer with a learning rate of 10^−3^ in both layers. DeepSpCas9 and DeepSpCas9-CA were implemented using TensorFlow (35).

### Training of conventional machine learning-based models

We trained seven models based on conventional machine learning algorithms, i.e., support vector machine (SVM), L1-regularized linear regression, L2-regularized linear regression, L1L2-regularized linear regression, AdaBoost, random forest, and gradient-boosted regression trees [ref]. All of the models were implemented using scikit-learn (ver 0.19.1)(36). A total of 627 features, which included position-independent and -dependent nucleotides and dinucleotides, melting temperature, GC counts, and the minimum self-folding free energy, were extracted as previously described (7,21). We performed 10-fold cross-validation for model selection among the regularization parameters and hyperparameter configurations, the number of which is comparable to the number of hyperparameter configurations used for the development of DeepSpCas9 (324). For L1-, L2-, and L1L2-regularized linear regression, over 250 points that were evenly spaced between 10^−6^ and 10^6^ in log space were searched to optimize the regularization parameter. For SVM, we searched over 225 models from the following hyperparameters: penalty parameter C and kernel parameter *γ*, 15 points that were evenly spaced between 10^−3^ and 10^3^. For random forest, AdaBoost, and gradient-boosted regression tree, we searched over 192 models selected from the following hyperparameter configurations: the number of base estimators (chosen from [50, 100, 200, 400]), the maximum depth of the individual regression estimators (chosen from [50, 100, 200, expanded until all leaves are pure]), the minimum number of samples to split an internal node (chosen from [2, 4]), the minimum number of samples to be at a leaf node (chosen from [1, 2]), and the maximum number of features to consider when looking for the best split (chosen from [all features, the square root of all features, the binary logarithm of all features]).

### Performance comparison of DeepSpCas9 with other models

We compared the prediction performance of DeepSpCas9 with those of the conventional machine learning-based models trained on the HT data set and other previously reported prediction models (7,10,13–16,18,37). The performance of each prediction model was evaluated by the Spearman correlation coefficients between experimentally measured sgRNA activities and prediction scores from each model. We used the Endo data set generated in this study and the other 14 published data sets from other groups that were large enough (number of target sequences > 100)(7,10,14,16,18,37–40) collected by Haeussler *et al*(20). In these test data sets, the target sequences included in the training data set HT were excluded. Furthermore, for a fair comparison of generalization performances, we excluded correlations of models tested against their own training data sets(20).

### Statistical significance

To compare the indel frequencies between DHS and non-DHS sites, we used the two-tailed Student t-test under the null hypothesis that the indel frequencies of the two groups are the same (Fig. 1b). To compare the Spearman correlation between prediction scores from two models (Fig. 2b, 2c, 2d, and 2e), we used Steiger’s test, which is used for testing two dependent correlation coefficients from exactly the same data set. Statistical significance was determined by using PASW Statistics (version 18.0, IBM) and Microsoft Excel (version, 16.0, Microsoft Corporation).

**Figure 1.**
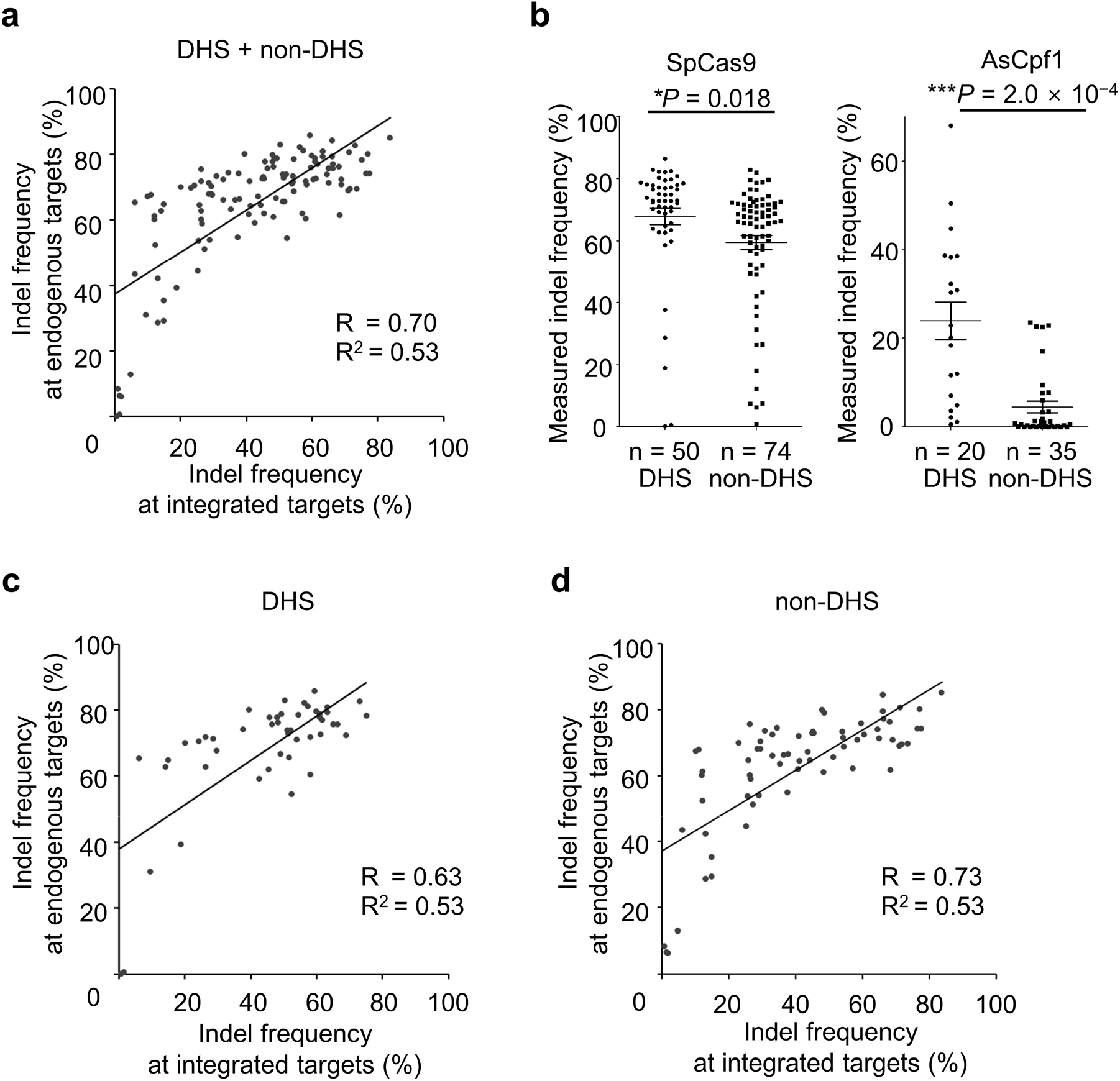
Correlation between indel frequencies measured by high-throughput approach and those by conventional method. **a** Correlation between indel frequencies at 120 endogenous and corresponding integrated target sequences. The Spearman correlation coefficients (R) and squared Pearson correlation coefficients (R^2^) are shown. **b** Effect of chromatin accessibility on the activities of SpCas9 (left) and AsCpf1 (right) at endogenous sites. Indel frequencies at endogenous sites were evaluated after transfection of plasmids encoding SpCas9 or AsCpf1 and guide RNAs. Indel frequencies at the target sites were compared after being divided into two groups, DNase I hypersensitive sites (DHS) and other sites (non-DHS). The numbers of analyzed target sites are as follows: (SpCas9) n = 62 for DHS target sites and n = 78 for non-DHS target sites, (AsCpf1) n = 20 for DHS target sites and n = 35 for non-DHS target sites. The HEK-plasmid data set from Ref. 20 was used for drawing this graph. Error bars represent s.e.m. Statistical significances determined by Student t-test are shown. **c**, **d** Correlation between indel frequencies at endogenous and corresponding integrated target sequences at 50 DNase I hypersensitive (DHS) sites (**c**) and 70 non-DHS sites (**d**). The Spearman correlation coefficients (R) and squared Pearson correlation coefficients (R^2^) are shown.

**Figure 2.**
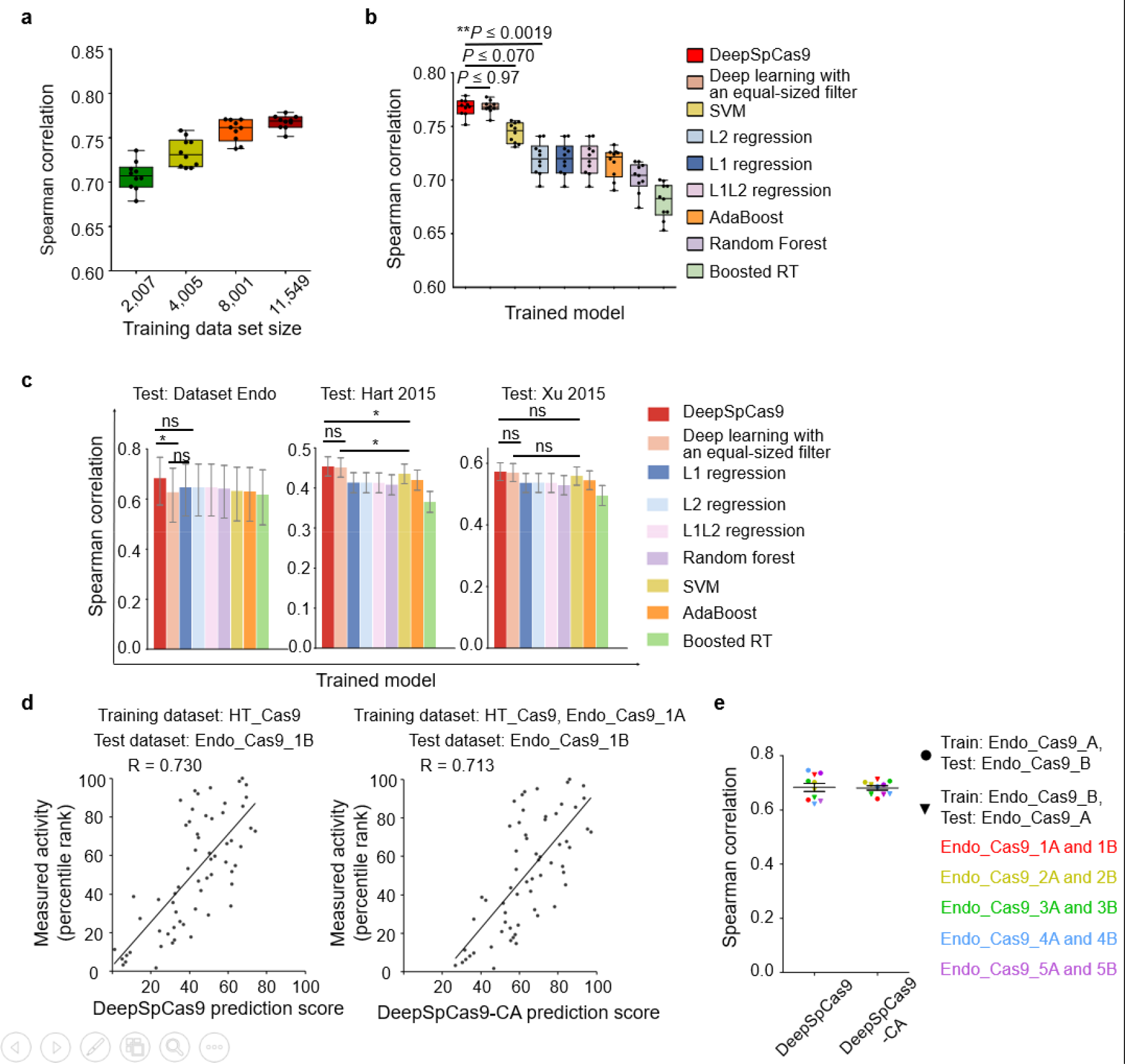
Evaluation of machine-learning based computational models predicting Cas9 activities. **a** Cross-validation of DeepSpCas9 models trained on data sets of varying sizes. Each dot represents the Spearman correlation coefficient between the measured indel frequency and the predicted activity from 10-fold cross-validation (total *n* = 10 correlation coefficients). In the boxes, the top, middle, and bottom lines represent the 25^th^, 50^th^, and 75^th^ percentiles, respectively, and whiskers indicate the minimum and maximum values. The confidence intervals are described in Supplementary Table 6. **b** Cross-validation of SpCas9 activity prediction models based on previously reported machine learning-based approaches. Each dot represents the Spearman correlation coefficient between the measured indel frequency and the predicted activity from 10-fold cross-validation (total *n* = 10 correlation coefficients). Statistical significances between the best, next-best, and third-best models are shown (DeepSpCas9 vs. deep learning with an equal-sized filter, *P* = 4.2 × 10^−2^, 2.9 × 10^−1^, 9.6 × 10^−1^, 9.7 × 10^−1^, 6.1 × 10^−1^, 9.0 × 10^−1^, 5.7 × 10^−1^, 2.9 × 10^−1^, 4.1 × 10^−1^, and 5.1 × 10^−1^ for each fold, respectively; DeepSpCas9 vs. SVM, *P* = 7.2 × 10^−3^, 7.5 × 10^−4^, 1.9 × 10^−3^, 1.1 × 10^−4^, 7.0 × 10^−2^, 5.9 × 10^−6^, 1.5 × 10^−2^, 2.0 × 10^−4^, 1.6 × 10^−3^, and 3.9 × 10^−2^ for each fold, respectively; DeepSpCas9 vs. L2 regression, *P* = 2.5 × 10^−7^, 8.3 × 10^−10^, 1.9 × 10^−8^, 2.5 × 10^−9^, 1.9 × 10^−3^, 2.1 × 10^−11^, 2.0 × 10^−7^, 1.4 × 10^−13^, 8.8 × 10^−10^, and 2.0 × 10^−4^ for each fold, respectively; Steiger’s test). In the boxes, the top, middle, and bottom lines represent the 25^th^, 50^th^, and 75^th^ percentiles, respectively. Whiskers indicate the minimum and maximum values. The confidence intervals are described in Supplementary Table 6. **c** Performance comparison of DeepSpCas9 with other prediction models using data set Endo_Cas9 (*n* = 124 independent target sites) and two published data sets (*n* = 4,207 and 2,060 independent target sites for data sets Hart 2015 and Xu 2015, respectively) as the test data sets. The bar graphs show the Spearman correlation coefficients between experimentally obtained indel frequencies and prediction scores for each of the models. Error bars represent 95% confidence intervals, which are described in detail in Supplementary Table 6. For clarity, results from statistical testing are shown only for DeepSpCas9 versus deep learning with an equal-sized filter, DeepSpCas9 versus the best conventional machine learning based model, and deep learning with an equal-sized filter versus the best conventional machine learning based model (left to right; \**P* = 1.4 × 10^−2^, DeepSpCas9 vs. deep learning with an equal-sized filter; \**P* = 1.1 × 10^−2^, DeepSpCas9 vs. SVM; \**P* = 4.6 × 10^−2^, deep learning with an equal size filter vs. SVM; ns = not significant; Steiger’s test). **d** Performance comparison of DeepSpCas9 and DeepSpCas9-CA (*c*hromatin *a*ccessibility). The DeepSpCas9-CA model was developed by fine-tuning the DeepSpCas9 model using the Endo-1A data set. DeepSpCas9 (left) and DeepSpCas9-CA (right) models were evaluated with the Endo-1B data set. The Spearman correlation coefficients (R) are shown. **e** Results from ten iterations of fine-tuning and evaluation. We divided the Endo_Cas9 data set into paired subsets by stratified random sampling from strata of DHS and nonDHS sites so that a similar ratio of DHS/nonDHS sites was assigned to each subset. We named the resulting data subset pairs Endo-1A and Endo-1B and repeated this sampling to generate four more pairs of data subsets, designated Endo-2A and Engo-2B, …, Endo-5A and Endo-5B. We next fine-tuned DeepSpCas9 using a data subset such as Endo-1A and the chromatin accessibility information, generating DeepSpCas9-CA. We compared the Spearman correlations of DeepSpCas9 and DeepSpCas9-CA using the other data subset, Endo-1B, as the test data. We next repeated this fine-tuning and subsequent testing after exchanging the training and test data sets: we used Endo-1B as the training data set for fine-tuning and Endo-1A as the test data set. We also conducted these analyses using the other four pairs of data sets. Each dot represents the Spearman correlation coefficient between the measured indel frequency and the predicted activity. A total of ten (= 2 × 5) rounds of fine-tuning and subsequent testing results are shown.

### Code availability

Source code for DeepSpCas9 and custom Python scripts used for the indel frequency calculations are available in Supplementary Code and on github (at https://github.com/MyungjaeSong/Paired-Library and https://github.com/CRISPRJWCHOI/IndelSearcher).

### Data availability

The deep sequencing data from this study have been submitted to the NCBI Sequence Read Archive (SRA; http://www.ncbi.nlm.nih.gov/sra/) under accession number SRP150719. The data sets used in this study are provided as Supplementary Table 1 and 2.

## RESULTS AND DISCUSSION

### Generation of large data sets of SpCas9 activities through a high-throughput evaluation

For a high-throughput evaluation of SpCas9 activities, we first prepared a lentiviral library of 15,656 guide RNA-encoding and target sequence pairs through a modification of our previous approach for Cpf1 (21,22). The target sequences were selected from the human genome and synthetic sequences without any information about the activity of the corresponding single-guide RNA (sgRNA) (detailed information is available in the Methods section). We prepared the paired plasmid library using two steps, i.e. Gibson assembly followed by restriction enzyme-induced cutting and ligation (Supplementary Fig. 1a), similar to the approach used for generating double guide RNA libraries (26,28,41–44) and libraries of guide RNA-encoding and target sequence pairs for the analysis of SpCas9-induced mutation patterns (23–25). Lentivirus was first generated from this plasmid library and then used to treat HEK293T cells to make a cell library, in which each cell contains a synthetic target sequence in its genome and expresses the corresponding sgRNA (Supplementary Fig. 1b). Next, the cell library was treated with SpCas9-encoding lentivirus, which led to sgRNA-directed cleavage and indel formation at the integrated target sequences with frequencies that depended on the sgRNA activity. The target sequences were PCR-amplified and subjected to deep sequencing to measure indel frequencies (21,22). This high-throughput experiment generated two data sets named HT_Cas9_Train and HT_Cas9_Test (Supplementary Tables 1 and 2).

### Indel frequencies at the integrated target sequences are highly correlated with those at the endogenous target sites

We next evaluated whether the indel frequencies at the integrated synthetic target sequences correlated with those at the corresponding endogenous target sites. For this determination, we generated a data set, named Endo_Cas9, of SpCas9 activities at 124 endogenous target sites with different chromatin accessibility properties (50 targets at DNase I hypersensitive (DHS) regions and 74 targets at non-DHS regions, Supplementary Table 3) because we previously found that Cpf1 activity is significantly affected by chromatin accessibility (22). We observed a strong correlation between indel frequencies at integrated target sequences and those at endogenous sites (Spearman R = 0.70, Pearson R^2^ = 0.53, Fig. 1a), which is higher than the correlation previously reported using a medium-scale library-on-library approach (18). Furthermore, we observed a weak tendency for SpCas9-induced indel frequencies at DHS sites to be marginally or merely higher than those at non-DHS sites (*P* = 0.018, Fig. 1b). In this respect, SpCas9 differs from Cpf1, which elicited dramatically higher levels of indels at DHS versus non-DHS sites (Fig. 1b) (22). When we calculated the correlations between indel frequencies at integrated sites and a subset of endogenous sites with similar chromatin accessibility, the correlations were comparable regardless of chromatin accessibility information (Fig. 1c,d). This observation also contrasts with previous observations of Cpf1, for which there was a much higher correlation between target site subsets with similar chromatin accessibility (22).

### Development of DeepSpCas9, a deep learning-based computational model predicting sgRNA efficacy

We next attempted to develop an accurate computational model for predicting SpCas9 activity. By using deep learning-based training on a large data set, we previously developed a computational model named DeepCpf1 that predicts AsCpf1 activity in a highly accurate manner (21). In this study, we used HT_Cas9_Train (Supplementary Table 1 and 2), a data set of SpCas9 activities at 12,832 integrated target sequences, which do not include target sequences used to generate Endo_Cas9 (Supplementary Table 2 and 3). By training on HT_Cas9_Train using an end-to-end deep-learning framework (31,34,45) (Supplementary Fig. 2), which is a modification of what we previously used to generate DeepCpf1, we developed DeepSpCas9, a deep learning-based regression model that predicts SpCas9 activity based on target sequence composition. Given that multiple filter sizes often improve the performance of deep learning(34), we used multiple sizes here instead of the equal-sized filters that we previously used(21). We conducted 10-fold cross-validation with HT_Cas9_Train to evaluate the generalization performance of model selection and training. As the size of the training data set for the cross-validation increased, the average Spearman correlation coefficients between experimentally obtained indel frequencies and predicted scores from DeepSpCas9 steadily increased up to 0.77 (Fig. 2a). When compared to conventional machine learning algorithms such as support vector machine (SVM), L1-regularized linear regression, L2-regularized linear regression, L1L2-regularized linear regression, AdaBoost, random forest, and gradient-boosted regression trees, which include those that previously showed competent performance for SpCas9 activity prediction (7,18), the Spearman correlations of DeepSpCas9 in the cross-validation were significantly higher than those of these conventional machine learning algorithms (vs. the second best model, SVM, *P* = 7.2 × 10^−3^, 7.5 × 10^−4^, 1.9 × 10^−3^, 1.1 × 10^−4^, 7.0 × 10^−2^, 5.9 × 10^−6^, 1.5 × 10^−2^, 2.0 × 10^−4^, 1.6 × 10^−3^, and 3.9 × 10^−2^ for each fold, respectively; vs. the third best model, L2-regression, *P* = 2.5 × 10^−7^, 8.3 × 10^−10^, 1.9 × 10^−8^, 2.5 × 10^−9^, 1.9 × 10^−3^, 2.1 × 10^−11^, 2.0 × 10^−7^, 1.4 × 10^−13^, 8.8 × 10^−10^, and 2.0 × 10^−4^ for each fold, respectively) and were similar to that of the equal-sized filter-based deep-learning model (Fig. 2b). Furthermore, when these algorithms were examined using the test data set Endo_Cas9 (derived using target sequences that were never included in the training data set HT_Cas9_Train) and two previously published data sets of Cas9 activities at endogenous sites (Hart 2015(38) and Xu 2015(16)), the Spearman correlation of DeepSpCas9 was also higher than those of the conventional machine-learning algorithms and that of the equal-sized filter-based deep-learning model (Fig. 2c), indicating that DeepSpCas9 exhibited the best performance among all of the models.

### Considering chromatin accessibility information barely improves SpCas9 activity prediction

We previously improved the prediction of Cpf1 activities at endogenous target sites by considering chromatin accessibility(21). To determine if such a consideration would also improve SpCas9 activity prediction, we divided the Endo_Cas9 data set into paired subsets by stratified random sampling from strata of DHS and nonDHS sites so that a similar ratio of DHS/nonDHS sites was assigned to each subset. We named the resulting data subset pairs Endo_Cas9_1A and Endo_Cas9_1B. We then repeated this stratified random sampling to generate four more pairs of data subsets, designated Endo_Cas9_2A and Endo_Cas9_2B, etc. (Supplementary Table 3). We next fine-tuned DeepSpCas9 using a data subset such as Endo_Cas9_1A and binary chromatin accessibility information from the Encyclopedia of DNA elements (ENCODE)(29), leading to the development of a fine-tuned model predicting SpCas9 activity based on both target sequence information and chromatin accessibility. When evaluated with the other data subset, Endo_Cas9_1B, as the test data set, the fine-tuned model showed performance comparable to that of DeepSpCas9 (Fig. 2d). We next repeated this fine-tuning and subsequent testing after exchanging the training and test data sets: we used Endo_Cas9_1B as the training data set for fine-tuning and Endo_Cas9_1A as the test data set. We also conducted these analyses using the other four pairs of data sets. This total of ten (= 2 × 5) rounds of fine-tuning and subsequent testing revealed that the Spearman correlations of these fine-tuned models are comparable to those of DeepSpCas9 (Fig. 2e), suggesting that fine-tuning with chromatin accessibility information barely improves the accuracy of DeepSpCas9 in predicting indel frequencies at endogenous sites. This result is in line with the finding that SpCas9 activity is only slightly affected by chromatin accessibility (Fig. 1b) and in strong contrast to DeepCpf1, which showed dramatically improved performance when chromatin accessibility information was considered (21).

### DeepSpCas9 shows unprecedentedly high generalization performance

To assess its generalization performance, we next evaluated DeepSpCas9 using other sufficiently large published data sets (number of target sequences > 100), derived from different studies from independent laboratories (seven data sets generated using U6 promoter-driven sgRNAs and three data sets generated using in vitro transcribed sgRNAs) (7,10,14,16,18,37–40), as test data and compared the results with those of other SpCas9-activity predicting programs (7,10,13–16,18,37). For a fair comparison of generalization performances, we excluded correlations of models tested against their own training data sets (20). We found that the Spearman correlations of DeepSpCas9 were the highest among those of nine previously published models in all seven tests against data sets generated using U6 promoter-driven sgRNAs and that statistical significance was observed for five out of the seven tests when compared with the second best models (Fig. 3), suggesting that DeepSpCas9 has the highest generalization performance compared to any of the other computational models predicting SpCas9 activity. DeepCRISPR(13), a recently reported deep-learning computational model trained using data sets of phenotypic changes of cells containing Cas9-induced gene edits, showed a lower generalization performance as compared to Doench 2016 (Rule set 2 or sgRNA designer)(7), which was developed before DeepCRISPR. When tested against the three data sets generated using in vitro transcribed sgRNA, the Spearman correlations of DeepSpCas9 were the highest together with those of CRISPRscan, which was generated for the prediction of in vitro transcribed sgRNA activities. Neither Doench 2016(7) nor CRISPRscan(10) showed the highest Spearman correlations for data sets of both U6 promoter-driven and in vitro transcribed sgRNAs. Taken together, these data suggest that the generalization performance of DeepSpCas9 is unprecedentedly high.

**Figure 3.**
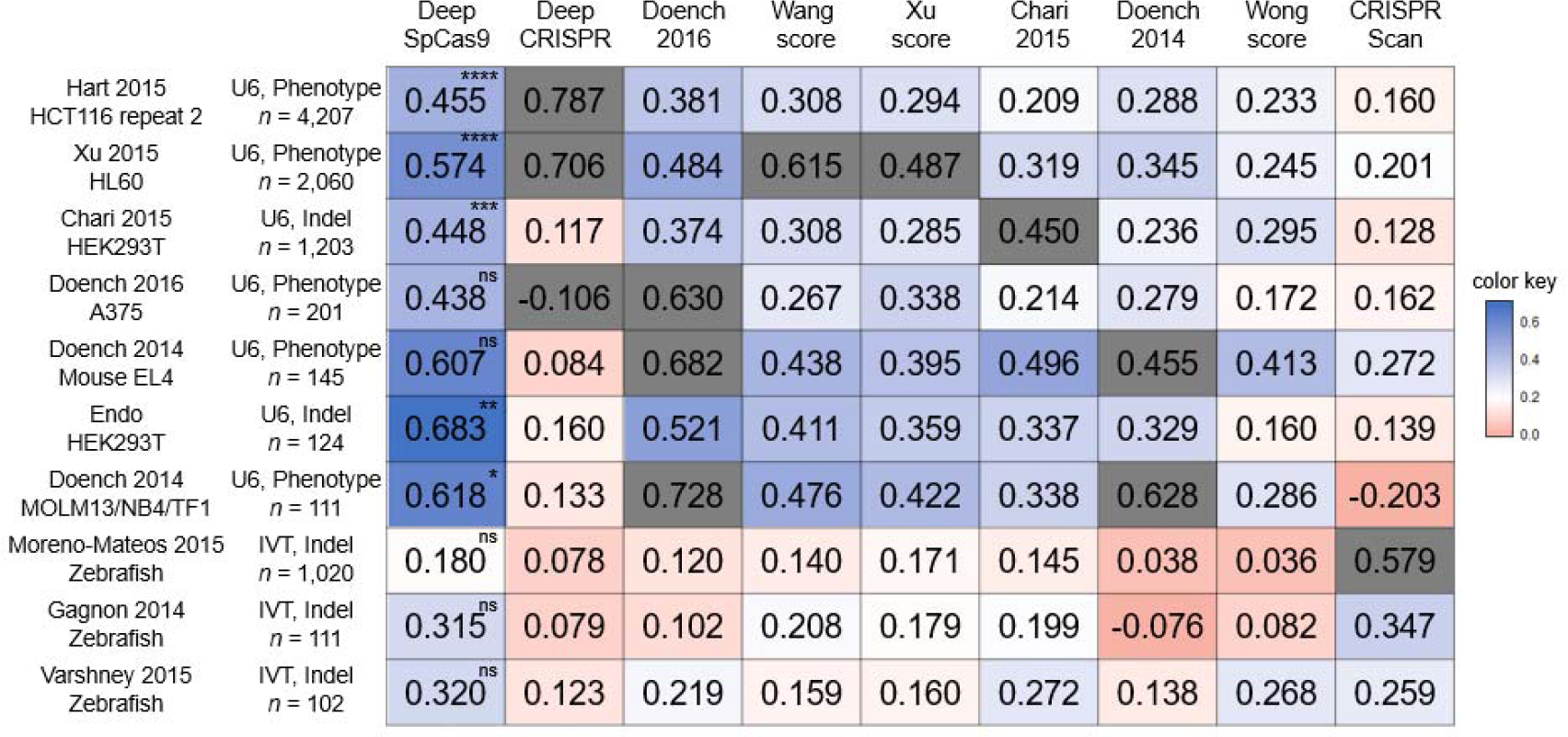
Comparison of generalization performances of computational models predicting Cas9 activities. The heat map shows Spearman correlation coefficients determined from DeepSpCas9 and previously reported models, which are arranged horizontally. The names of the vertically placed test data sets include information about the cell line or species used. Other related parameters, such as the guide RNA expression method (U6 promoter-driven (U6) vs. in vitro transcribed (IVT)), the Cas9 activity analysis method (phenotypic change (phenotype) vs. indel), and the number of analyzed sites, are also shown. Each grey box indicates the correlation of a model tested against a test data set that includes its own training data set. In the evaluation against each test data set, the statistical significance between the two best models is indicated for the best model (From the top; \*\*\*\**P* = 5.3 × 10^−9^, \*\*\*\**P* = 1.8 × 10^−10^, \*\*\*\**P* = 3.4 × 10^−8^, \*\*\*\**P* = 1.1 × 10^−13^, \*\*\*\**P* = 2.9 × 10^−11^, \*\*\*\**P* = 3.9 × 10^−8^, \*\*\**P* = 2.5 × 10^−4^, ns = not significant, \**P* = 3.7 × 10^−2^, and \**P* = 3.9 × 10^−2^; Steiger’s test).

We provide a web tool that enables accurate prediction of SpCas9 activity by DeepSpCas9 at http://deepcrispr.info/DeepCas9 and provide code for incorporation of DeepSpCas9 into existing tools (Supplementary Code). Given that DeepSpCas9 has unprecedentedly high generalization performance, we expect that it will greatly facilitate genome editing using SpCas9.

## Author contributions

H.K.K. and Y.K. performed experiments to build high-throughput data sets of SpCas9 indel frequencies. H.K.K., Y.K., and J.Y.B. performed experiments to build data sets of SpCas9 indel frequencies at endogenous sites. S.L., S.M., and S.Y. developed the framework, and carried out the model training and computational validation. J.W.C., J.P, and D.J. significantly contributed to bioinformatics analyses. H.K. conceived and designed the study. H.K.K., Y.K., S.L., S.M., S.Y., and H.K. analyzed the data and wrote the manuscript.

## Acknowledgments

This work was supported by the Korean Health Technology R&D Project, Ministry of Health and Welfare, Republic of Korea (HI17C0676 (H.K.)).

## Supplementary Information

**Fig. S1.**
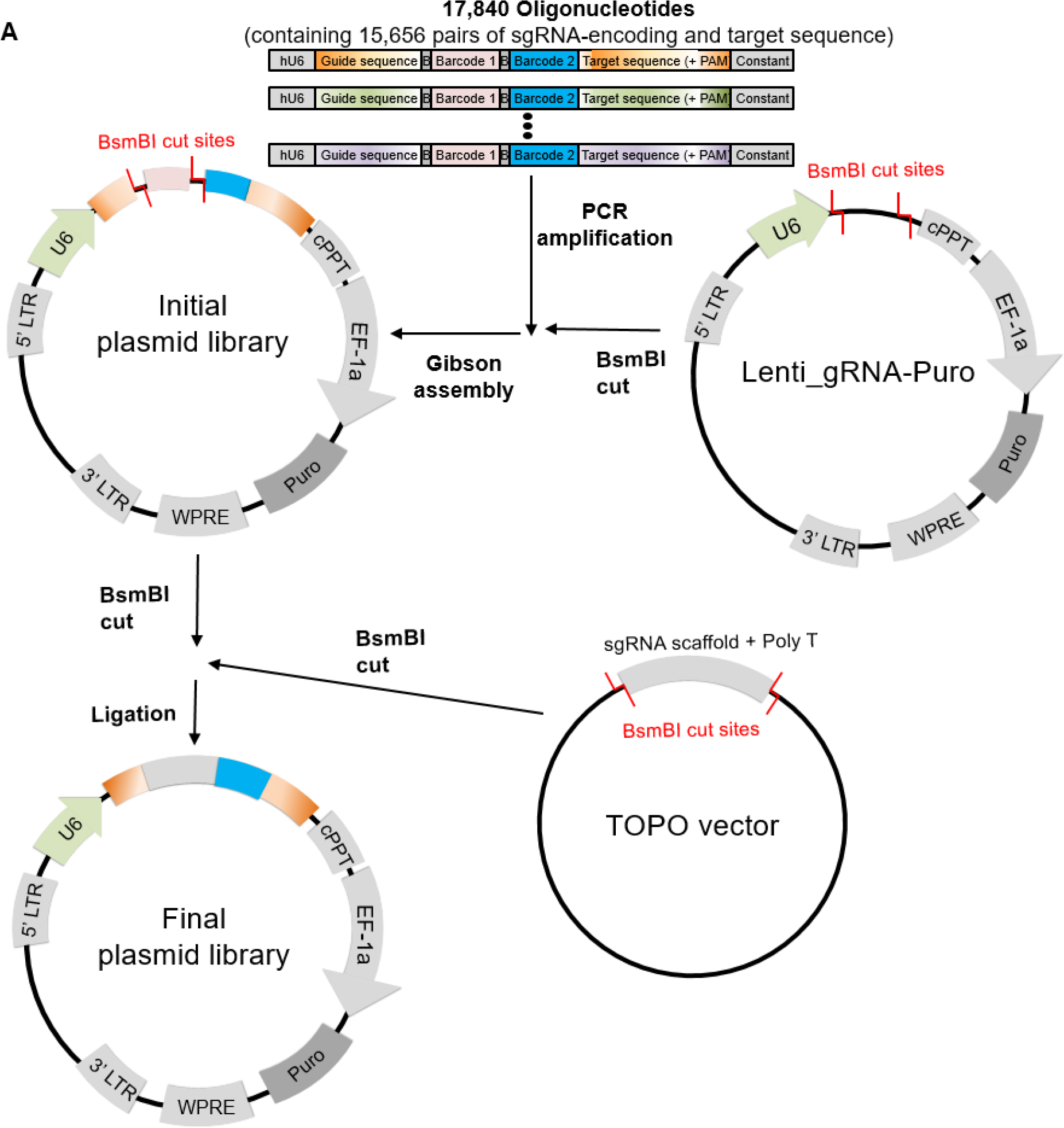
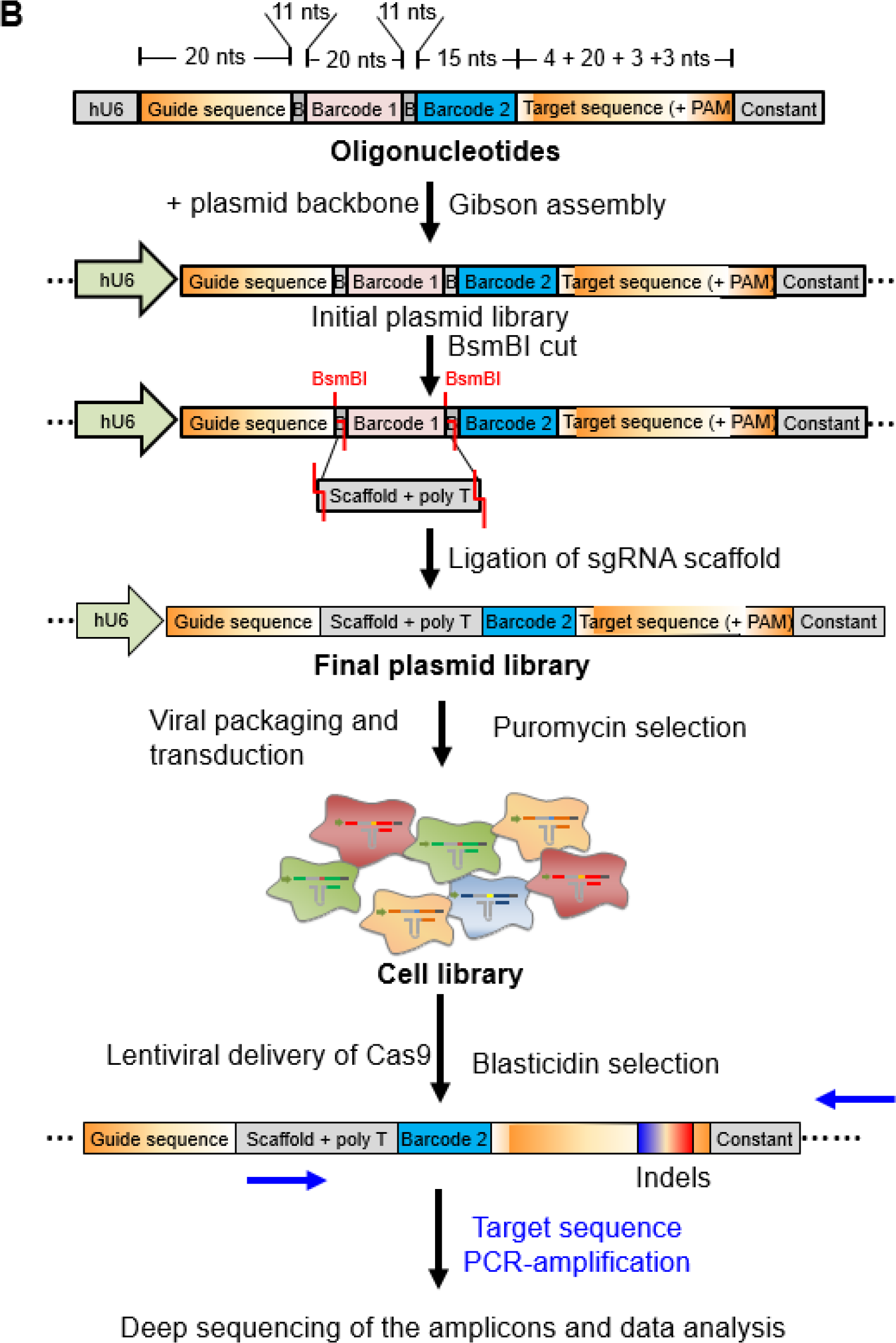
Development of a high-throughput evaluation system for Cas9-induced indel frequencies. **(A)** Overview of the cloning strategy for generating the plasmid library of sgRNA-encoding and target sequence pairs. A pool of 17,840 oligonucleotides, each containing a guide RNA-target sequence pair, was PCR-amplified and cloned into the Lenti_gRNA-Puro vector using Gibson assembly. This initial plasmid library was linearized using BsmBI digestion and ligated with a BsmBI-digested guide RNA scaffold fragment to generate the final plasmid library. (**B**) A schematic of the high-throughput evaluation system used in this study. The pool of oligonucleotides, each containing a 20-nt sgRNA guide sequence, a BsmBI restriction site, a 20-nt barcode sequence (barcode 1), a second BsmBI restriction site, a 15-nt barcode sequence (barcode 2), and the 30-nt corresponding target sequence including a PAM, were PCR-amplified and cloned into a plasmid using Gibson assembly. The sgRNA scaffold sequence was inserted into this initial plasmid library using BsmBI-induced cutting and subsequent ligation. The resulting final plasmid library was used to generate a lentiviral library, which was in turn used to treat HEK293T cells to create a cell library. Lentiviral delivery of Cas9 into this cell library induced indels at the integrated target sequences with frequencies that depended on the sgRNA activity.

**Fig. S2.**
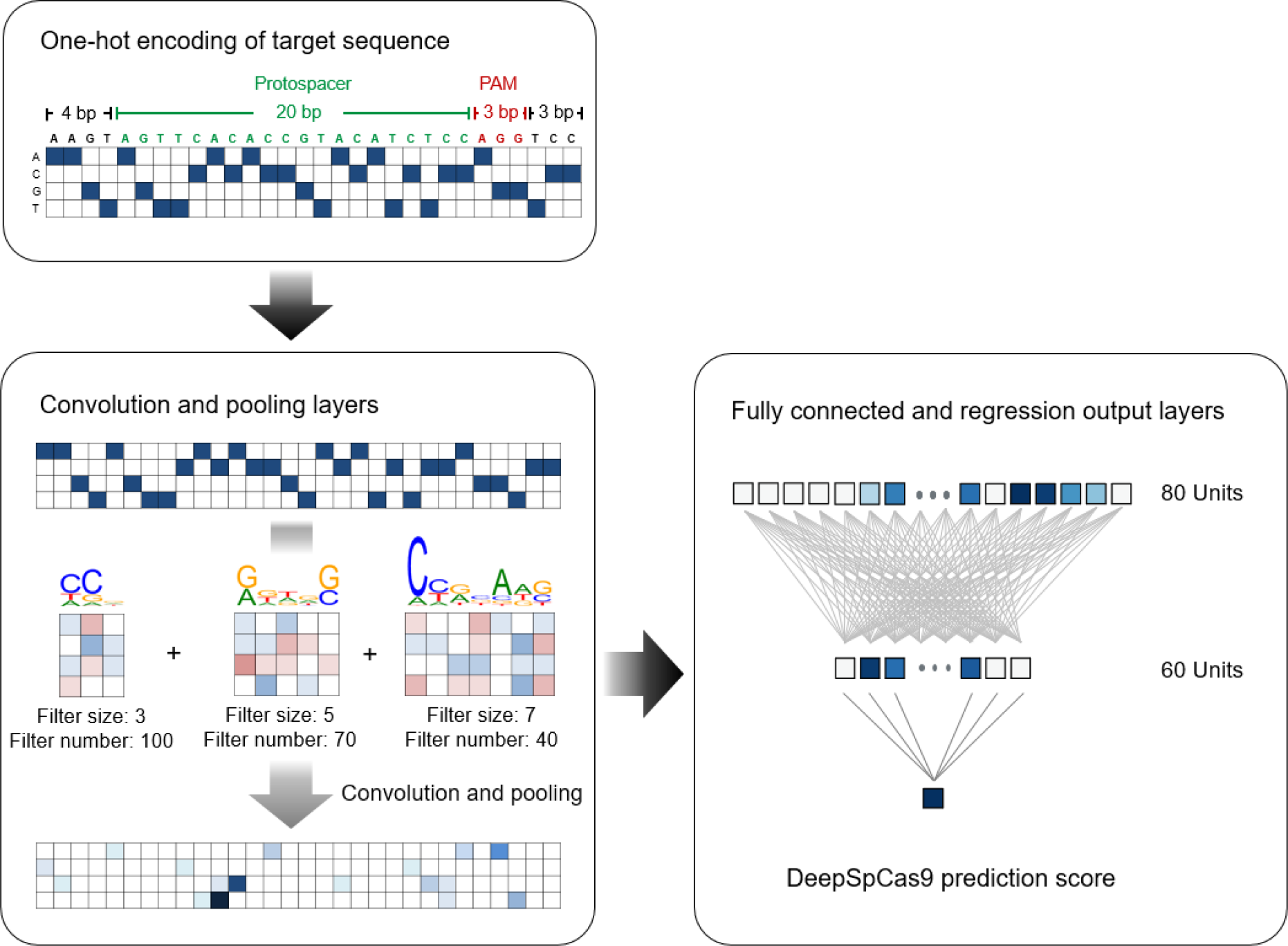
Overview of DeepSpCas9 development. DeepSpCas9 is based on a deep learning framework using robust convolutional neural networks (CNNs). DeepCas9 development involved the following steps. (1) 30 base pair (bp) input sequences that include the target and neighboring sequences were converted into a four-dimensional binary matrix. (2) A total of 210 filters (100, 70, and 40 filters at 3, 5, and 7 nts in length, respectively) were shifted through the four-dimensional binary matrix to determine the position weight matrices. The maximum values were pooled from the local features calculated from the previous convolution layer to “pool out” those that were informative. (3) Pooled features were then combined according to the weighted sum and rectified linear unit non-linear function in the fully connected layers. (4) The output layer performs linear regression and predicts the activity score for each SpCas9 guide RNA.

**Table S1**. Data sets generated from the results of the high-throughput experiments (provided in separate file)

**Table S2.**
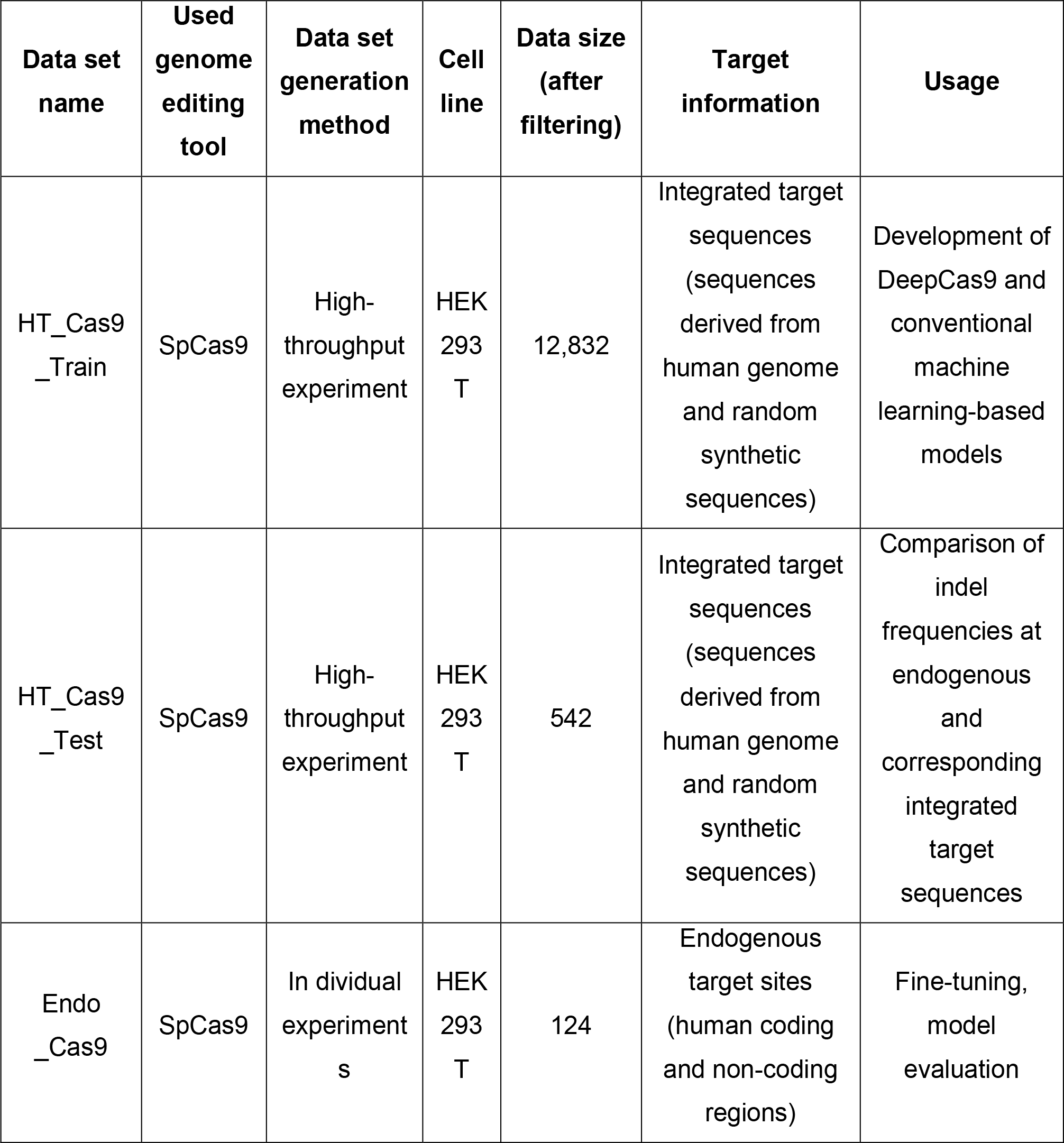
Datasets used for this study. Datasets HT_Cas9_Train, HT_Cas9_Test contain the indel frequencies at integrated target sequences and were obtained from independent high-throughput experiments conducted in HEK293T cells. Indel frequencies at endogenous human coding and non-coding regions were included in dataset Endo_Cas9.

**Table S3**. Data sets generated from the results of the experiments at endogenous target sites. (provided in separate excel file)

**Table S4.** Oligonucleotides used in this study. (provided in separate excel file)

**Table S5**. Model selection results (provided as a separate excel file). Of the 324 models that were explored, model 54 was selected to define the final model for DeepCas9 and is highlighted in red.

**Table S6**. Confidence intervals for the values shown in the graphs. (provided in separate file)

