## Supplementary table 2 for "SpCas9 activity prediction by DeepSpCas9, a deep learning-based model with unparalleled generalization performance"

**Supplementary Table 2.** Datasets used for this study. Datasets HT_Cas9_Train, HT_Cas9_Test contain the indel frequencies at integrated target sequences and were obtained from independent high-throughput experiments conducted in HEK293T cells. Indel frequencies at endogenous human coding and non-coding regions were included in dataset Endo_Cas9.

| **Data set name** | **Used genome editing tool** | **Data set generation method** | **Cell line** | **Data size (after filtering)** | **Target information** | **Usage** |
|---|---|---|---|---|---|---|
| HT_Cas9 _Train | SpCas9 | High-throughput experiment | HEK 293 T | 12,832 | Integrated target sequences (sequences derived from human genome and random synthetic sequences) | Development of DeepCas9 and conventional machine learning-based models |
| HT_Cas9 _Test | SpCas9 | High-throughput experiment | HEK 293 T | 542 | Integrated target sequences (sequences derived from human genome and random synthetic sequences) | Comparison of indel frequencies at endogenous and corresponding integrated target sequences |
| Endo _Cas9 | SpCas9 | In dividual experiment s | HEK 293 T | 124 | Endogenous target sites (human coding and non-coding regions) | Fine-tuning, model evaluation |
